## Supplementary Information for "Trainable subnetworks reveal insights into structure knowledge organization in protein language models"

### S1 Supplementary Text

#### S1.1 Subnetwork training details

For each PLM, we conducted a grid search to select optimal hyperparameter values to achieve suppression in all subnetworks. We set the below hyperparameters to the same values across all subnetworks learned within the same model. We include the results of the grid search on subnetwork performance in the accompanying code and data (see https://github.com/microsoft/plm_subnetworks for more information). The hyperparameters that vary by PLM are listed below and summarized in Table S1.

- **Mask initialization ($s_{init}$).** Each mask logit $l_i$ is initialized such that $\sigma(l_i) = p_{\text{keep},0}$, where $p_{\text{keep},0}$ is the desired initial retention probability (e.g., 0.96) of the $i$-th parameter. This is achieved by setting $l_i = \log\left[p_{\text{keep},0}/(1 - p_{\text{keep},0})\right]$, so that each parameter begins training with a high probability of being retained. This initialization enables the model to start from a near-complete network and learn to suppress parameters selectively over time. We choose $p_{\text{keep},0}$ to ensure that the subnetwork MLM loss initially matches the PLM average loss over all inputs, while introducing less than approximately 1% sparsity. This yields a mask score distribution centered near zero, placing most sigmoid activations in a high-sensitivity regime and allowing flexible updates toward maintenance and suppression goals. We explored the effects of this random initialization of mask scores in Fig. S2.
- **Suppression-KL loss weight ($\lambda_{\text{supp.}}$).** Applied using Eq. (1) and Eq. (2).
- **Maintenance-KL loss weight ($\lambda_{\text{maint.}}$).** Applied using Eq. (3) and Eq. (4).
- **Maintenance-MLM loss weight ($\lambda_{\text{MLM}}$).** Applied using Eq. (5) and Eq. (6).
- **Mask temperature ($\tau$).** We apply a cosine decay schedule to the temperature $\tau$, decreasing it smoothly from a maximum of $\tau_{\text{init}}$ to $\tau_{\text{final}}$ over 100 epochs. This schedule enables high-entropy exploration early in training and progressively sharper binarization of the mask as training progresses.
- **Mask threshold ($T$).** In practice, the binarization threshold $T$ in Eq. (9) varies by PLM.

#### S1.2 Learning rate

We use a maximum learning rate of `peak_lr` with a reverse OneCycle learning rate schedule. The learning rate remains constant at `peak_lr` during an initial plateau phase, defined by `1 / lr_plateau_epochs`, and then decays to `peak_lr` divided by 30 using cosine annealing. The learning rate schedule spans the full number of estimated training steps, as determined by PyTorch Lightning. Only mask parameters are optimized, using the AdamW optimizer with $\beta = (0.9, 0.999)$, $\epsilon = 10^{-6}$, and zero weight decay. The maximum value of `peak_lr` observed in training is $10^{-1}$.

#### S1.3 Compute

We train each subnetwork on one NVIDIA H100 GPU. We curate training batches of size 16 with a 0.25 frequency of suppression inputs in a batch such that subnetworks training converges over fewer epochs. We report the number of suppression inputs per category and the number training steps in Table S3.

### Supplementary Figures

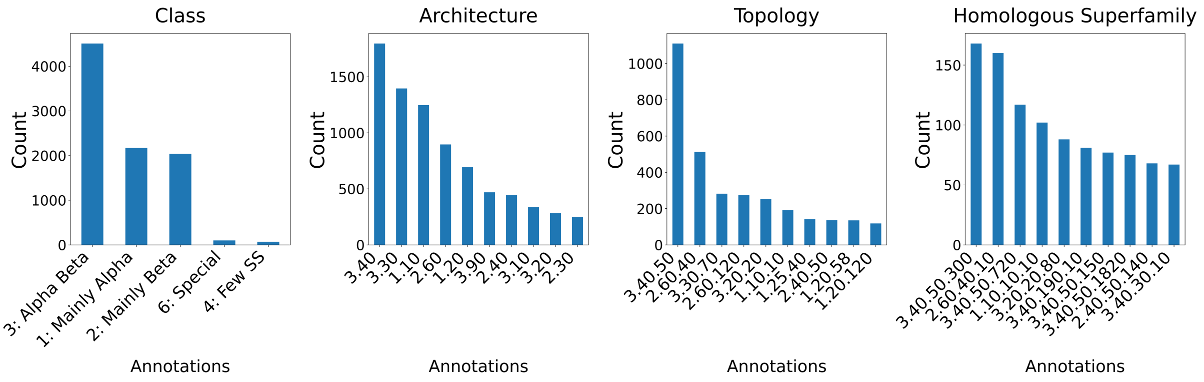


**(A) Annotation frequencies by CATH levels.** Each CATH domain is annotated with a label at the Class, Architecture, Topology, and Homologous Superfamily levels. Bar plots show the counts (y-axis) of the top 10 most frequent annotations (x-axis) at each level of the CATH hierarchy.

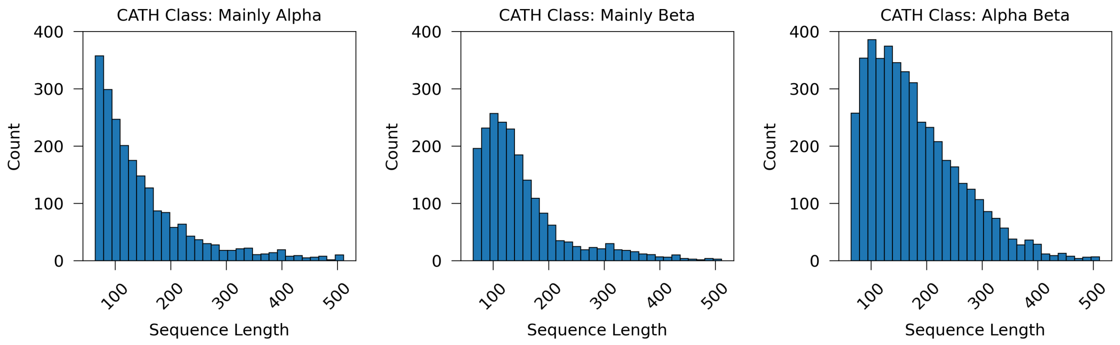


**(B) Sequence length distributions stratified by CATH level.** Histograms show the counts (y-axis) of domain sequence lengths (x-axis) for each CATH Class: Mainly Alpha, Mainly Beta, Alpha Beta.

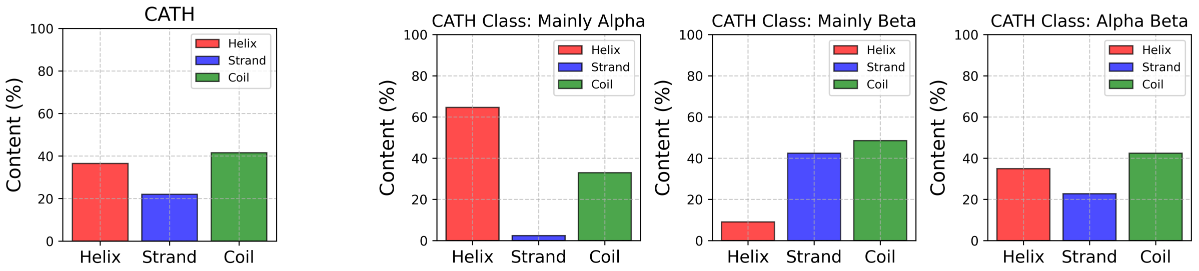


**(C) Secondary structure composition of CATH domains.** (Left) Average fraction of residues annotated as helix, strand, or coil across all CATH domains, based on DSSP annotations [28]. (Right) Breakdown of secondary structure content stratified by CATH class. DSSP 8-state annotations are mapped to 3-state labels: H, G, I → H; E, B → E; T, S, - → L. Helix is H, beta strand is E, and loop is L.

**Figure S1. CATH annotation characteristics.**

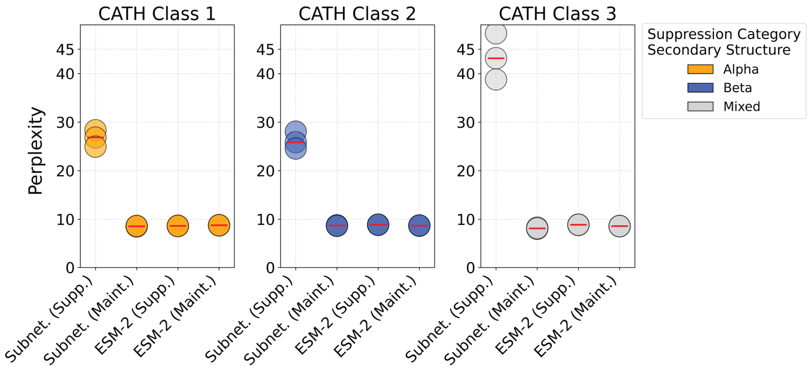


**Figure S2. ESM-2 CATH Class-level subnetwork language modeling performance across multiple seeds.** Three subnetworks were independently trained for each CATH Class suppression target (Mainly Alpha, Mainly Beta, Alpha-Beta) to assess the reproducibility of mask learning given random initialization of mask scores. Each point represents the validation perplexity of a subnetwork stratified by category of inputs.

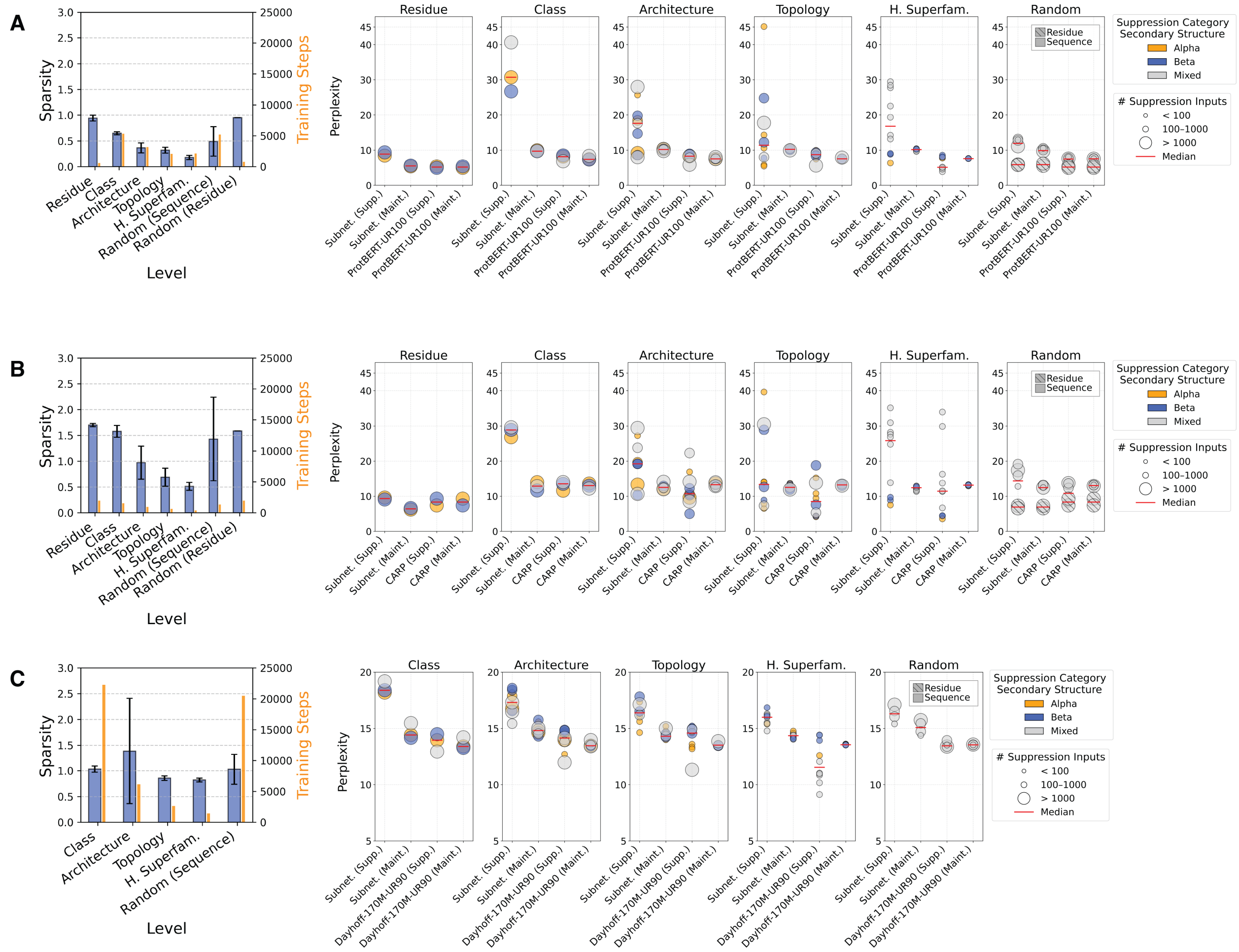


**Figure S3. Subnetwork language modeling performance on three additional PLMs of varying size and architectures.** To investigate whether subnetworks exist in pretrained PLMs other than ESM-2, we applied our approach to three additional models of varying size and architectures trained on UniRef data: ProtBERT-UR100, a transformer masked language model with a BERT architecture [2]; CARP-640M, a convolutional neural network masked language model (transformer layers are replaced by ByteNet dilated CNN blocks) [14]; and Dayhoff-170M-UR90, an efficient hybrid state-space-model transformer trained with and autoregressive objective [31]. **(A) ProtBERT-UR100 (420M).** Left: Learned percent sparsity by category of learned subnetworks in ProtBERT-UR100. Right: Masked language modeling performance on suppression and maintenance categories of sequences for each subnetwork. **(B) CARP-640M.** Left: Learned percent sparsity by category of learned subnetworks in CARP-640M. Right: Masked language modeling performance on suppression and maintenance categories of sequences for each subnetwork. **(C) Dayhoff-170M-UR90.** Left: Learned percent sparsity by category of learned subnetworks in Dayhoff-170M-UR90. Right: Autoregressive language modeling performance on suppression and maintenance categories of sequences for each subnetwork. Residue subnetworks were omitted from Dayhoff-170M-UR90 results because causal perplexity cannot be computed selectively over individual residues.

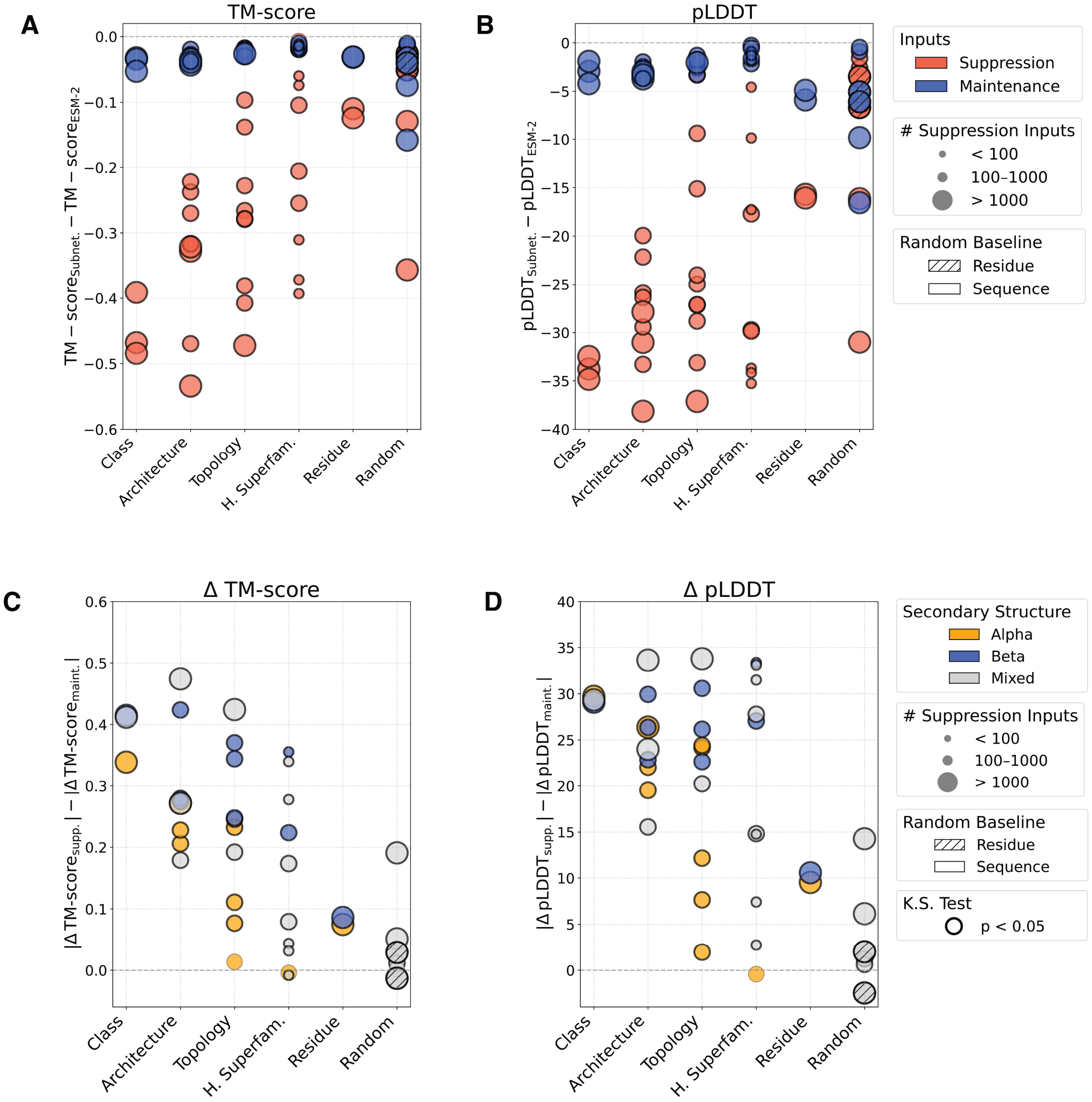


**Figure S4. ESM-2 650M subnetwork-predicted TM-score and pLDDT with the ESM-2 folding trunk.** **(A–B)** Structural prediction differences (y-axis) between subnetworks and the ESM-2 baseline shown for **(A)** TM-score and **(B)** pLDDT across structural levels (x-axis). Each point represents change in metrics for suppression inputs (red) or maintenance inputs (blue) for a subnetwork relative to ESM-2. Marker size indicates the number of suppression inputs for each subnetwork. Bold outlines of markers indicate statistically significant paired t-test $p$-values ($p < 0.05$). **(C–D)** Difference in absolute structure prediction metric changes from the ESM-2 baseline (y-axis), stratified by structural level, for **(C)** TM-score and **(D)** pLDDT. Each point corresponds to an individual subnetwork and shows the difference between suppression and maintenance Δ-values. Marker size reflects the number of suppressed inputs in the subnetwork, and color indicates secondary structure type. Bold outlines of markers indicate statistical significance by Kolmogorov–Smirnov (KS) test ($p < 0.05$).

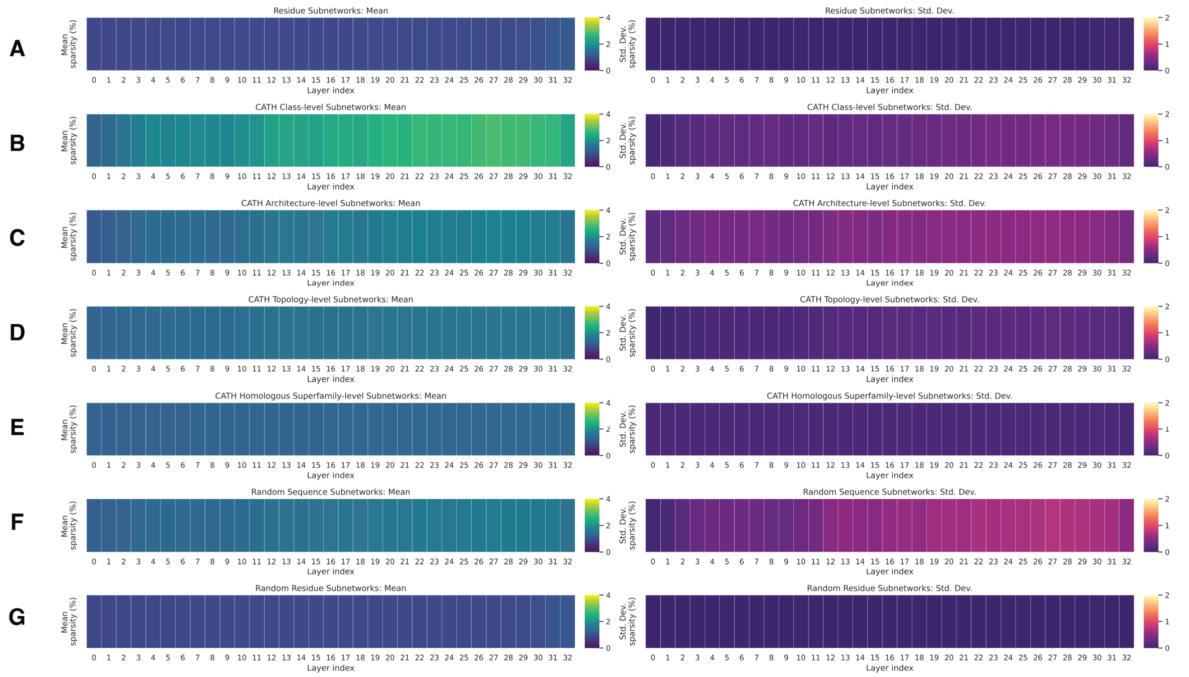


**Figure S5. Mask interpretation of ESM-2 650M.** Mean and standard deviation percent of parameters pruned by layer for subnetworks grouped at the levels of **(A)** residue, **(B)** CATH class, **(C)** CATH architecture, **(D)** CATH topology, **(E)** CATH homologous superfamily, **(F)** random sequence suppression, and **(G)** random residue suppression.

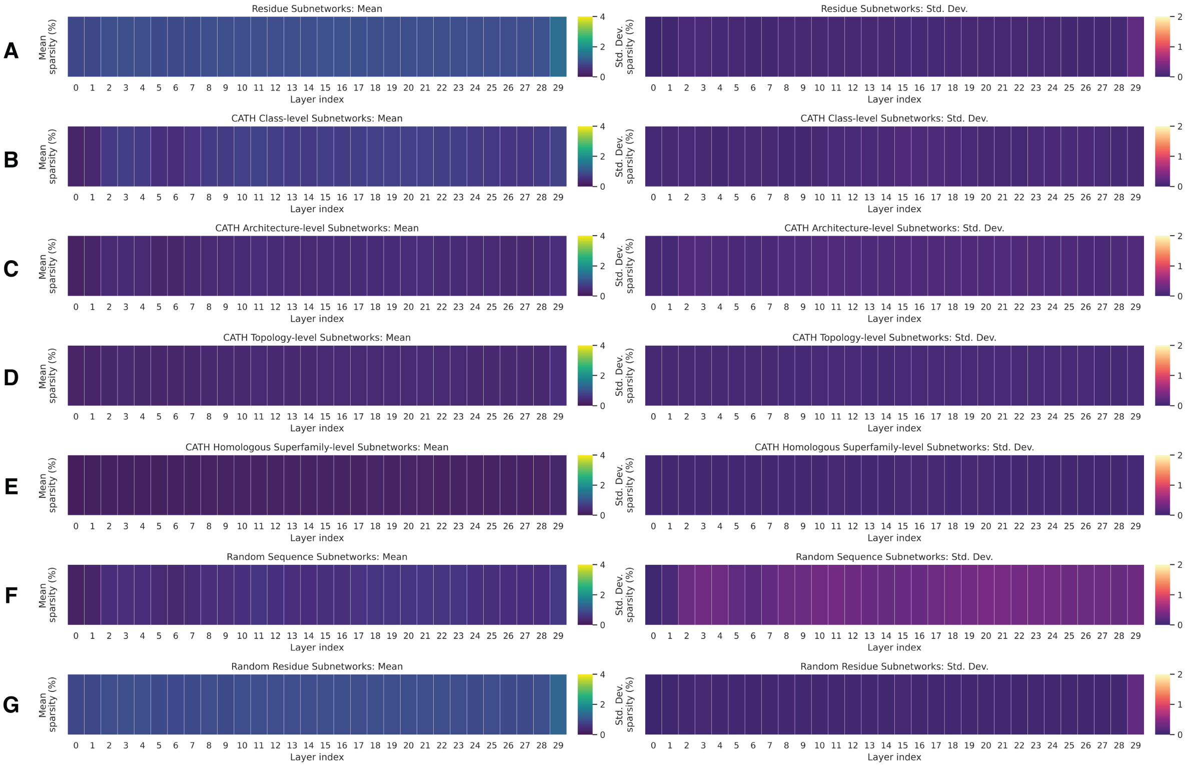


**Figure S6. Mask interpretation of ProtBERT-UR100.** Mean and standard deviation percent of parameters pruned by layer for subnetworks grouped at the levels of **(A)** residue, **(B)** CATH class, **(C)** CATH architecture, **(D)** CATH topology, **(E)** CATH homologous superfamily, **(F)** random sequence suppression, and **(G)** random residue suppression.

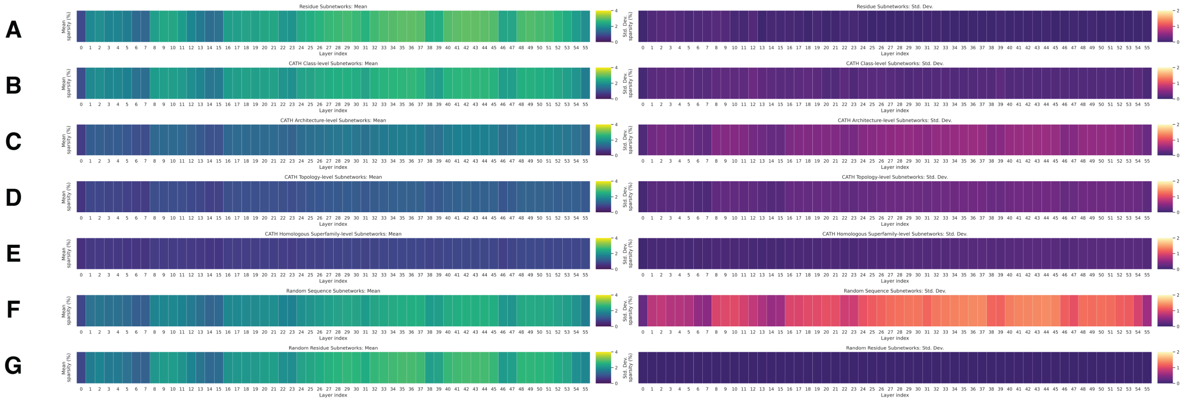


**Figure S7. Mask interpretation of CARP-640M.** Mean and standard deviation percent of parameters pruned by layer for subnetworks grouped at the levels of **(A)** residue, **(B)** CATH class, **(C)** CATH architecture, **(D)** CATH topology, **(E)** CATH homologous superfamily, **(F)** random sequence suppression, and **(G)** random residue suppression.

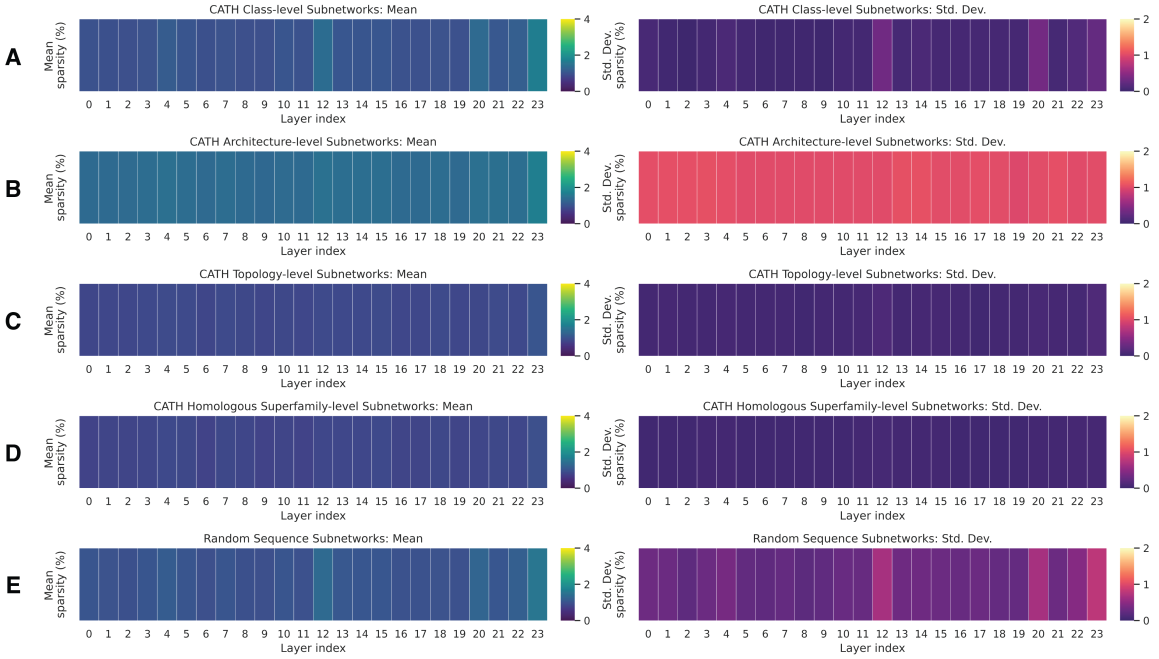


**Figure S8. Mask interpretation of Dayhoff-170M-UR90.** Mean and standard deviation percent of parameters pruned by layer for subnetworks grouped at the levels of **(A)** CATH class, **(B)** CATH architecture, **(C)** CATH topology, **(D)** CATH homologous superfamily, and **(E)** random sequence suppression.

### Supplementary Tables

**Table S1. Training and hyperparamter configurations and masked modules for subnetwork learning across four pretrained protein language models.** Each PLM subnetwork differs in which modules are masked, according to the model architecture, following evidence that knowledge is localized in representational modules [22]. Masking is thus applied to self-attention or convolutional projections while leaving embeddings, bias, and normalization layers intact. Subnetworks trained within a single PLM reuse the same hyperparameter configuration shown below. ESM-2 650M, ProtBERT-UR100, and CARP-640M are masked-language-model PLMs trained with the same MLM objective, whereas Dayhoff-170M-UR90 is an autoregressive model trained with a next-token prediction objective.

| Model | Masked Modules | Layers | $s_{\text{init}}$ | $\lambda_{\text{maint}}$ | $\lambda_{\text{supp}}$ | $\lambda_{\text{MLM}}$ | $\tau_{\text{init}}$ | $\tau_{\text{final}}$ | $T$ |
|---|---|---|---|---|---|---|---|---|---|
| ESM-2 650M | Self-attention layers | 34 | 0.96 | 7 | 10 | 1 | 3 | 0.01 | 0.40 |
| ProtBERT-UR100 | Self-attention layers | 30 | 0.998 | 9 | 10 | 1 | 1 | 0.1 | 0.43 |
| CARP-640M | Convolutional kernels | 56 | 0.995 | 9 | 10 | 1 | 1 | 0.1 | 0.40 |
| Dayhoff-170M-UR90 | Self-attention + Mamba projections | 24 | 0.99 | 4 | 4 | 4 | 1 | 0.1 | 0.40 |

**Table S2. Mean and standard deviation of ESM-2 CATH Class-level subnetwork masked language modeling performance across multiple seeds.** Three subnetworks were independently trained for each CATH Class suppression target (Mainly Alpha, Mainly Beta, Alpha-Beta) to assess the reproducibility of mask learning given random initialization of mask scores. Reported are the mean ± standard deviation of masked language modeling perplexity for subnetworks and baseline ESM-2 perplexity stratified by categories of inputs.

| **Category** | **Target** | **Sparsity** | **Training Step** | **Subnet. Supp.** | **Subnet. Maint.** | **ESM Supp.** | **ESM Maint.** | **t-Test ($p$) Supp.** | **t-Test ($p$) Maint.** |
|---|---|---|---|---|---|---|---|---|---|
| Class | 1 | 2.47 | 7,600 | $24.9 \pm 72.5$ | $8.6 \pm 17.4$ | $8.6 \pm 15.2$ | $8.8 \pm 14.1$ | $< 1e{-16}$ | 3.7e-05 |
| | 1 | 2.45 | 7,640 | $26.8 \pm 82.5$ | $8.6 \pm 16.1$ | $8.6 \pm 15.2$ | $8.8 \pm 14.1$ | $< 1e{-16}$ | 2.4e-12 |
| | 1 | 2.48 | 7,560 | $28.3 \pm 82.9$ | $8.6 \pm 16.2$ | $8.6 \pm 15.2$ | $8.8 \pm 14.1$ | $< 1e{-16}$ | 2.2e-13 |
| | 2 | 2.4 | 7,040 | $28.0 \pm 70.2$ | $8.7 \pm 16.0$ | $8.9 \pm 15.7$ | $8.7 \pm 14.0$ | $< 1e{-16}$ | 6.5e-03 |
| | 2 | 2.36 | 7,160 | $25.8 \pm 62.8$ | $8.7 \pm 16.3$ | $8.9 \pm 15.7$ | $8.7 \pm 14.0$ | $< 1e{-16}$ | 1.5e-02 |
| | 2 | 2.4 | 7,120 | $24.6 \pm 59.8$ | $8.7 \pm 15.4$ | $8.9 \pm 15.7$ | $8.7 \pm 14.0$ | $< 1e{-16}$ | 3.0e-01 |
| | 3 | 2.56 | 7,880 | $48.3 \pm 106.4$ | $8.3 \pm 22.4$ | $8.9 \pm 13.7$ | $8.6 \pm 15.2$ | $< 1e{-16}$ | $< 1e{-16}$ |
| | 3 | 2.58 | 7,900 | $43.2 \pm 89.9$ | $8.1 \pm 17.0$ | $8.9 \pm 13.7$ | $8.6 \pm 15.2$ | $< 1e{-16}$ | $< 1e{-16}$ |
| | 3 | 2.59 | 8,000 | $38.8 \pm 84.3$ | $8.1 \pm 18.0$ | $8.9 \pm 13.7$ | $8.6 \pm 15.2$ | $< 1e{-16}$ | $< 1e{-16}$ |

**Table S3. Overview of subnetwork definitions.** Each subnetwork is trained to selectively suppress either a specific type of residue belonging to a secondary structure or a set of sequences that belong to the same CATH category classification. In our study, we consider only the top 10 most frequent labels in each CATH category, and report the number of sequences and predominant type of secondary structure content in each category.

| **Category** | **Target** | **# Seqs.** | **Type** |
|---|---|---|---|
| Residue | Helix | 36.5% of residues | Alpha |
| | Sheet | 21.9% of residues | Beta |
| Class | 1 | 2169 | Alpha |

| Category | Target | # Seqs. | Type |
|---|---|---|---|
| | 2 | 2038 | Beta |
| | 3 | 3129 | Mixed |
| Architecture | 1.10 | 1247 | Alpha |
| | 1.25 | 182 | Alpha |
| | 1.20 | 693 | Alpha |
| | 2.30 | 251 | Beta |
| | 2.40 | 447 | Beta |
| | 2.60 | 896 | Beta |
| | 3.30 | 1396 | Mixed |
| | 3.40 | 1797 | Mixed |
| | 40 | 469 | Mixed |
| Topology | 1.10.10 | 192 | Alpha |
| | 1.10.287 | 111 | Alpha |
| | 1.20.58 | 135 | Alpha |
| | 1.20.120 | 118 | Alpha |
| | 1.25.40 | 142 | Alpha |
| | 2.40.50 | 136 | Beta |
| | 2.60.40 | 512 | Beta |
| | 2.60.120 | 276 | Beta |
| | 3.30.70 | 282 | Mixed |
| | 3.40.50 | 1110 | Mixed |
| H. Superfam. | 3.40.50.300 | 168 | Mixed |
| | 2.60.40.10 | 160 | Beta |
| | 3.40.50.720 | 117 | Mixed |
| | 1.10.10.10 | 102 | Alpha |
| | 3.20.20.80 | 88 | Mixed |
| | 3.40.190.10 | 81 | Mixed |
| | 3.40.50.150 | 77 | Mixed |
| | 3.40.50.1820 | 75 | Mixed |
| | 2.40.50.140 | 68 | Beta |
| | 3.40.30.10 | 67 | Mixed |
| Random | Sequences | 100 | Mixed |
| | | 200 | Mixed |
| | | 1000 | Mixed |
| | | 2000 | Mixed |
| Random | Residue | Random | Mixed |

**Table S4. ESM-2 650M subnetwork and baseline language modeling performance.** Below we report the mean and standard deviation of the subnetwork and ESM-2 baseline perplexities (illustrated in Fig. 2C) stratified by categories of inputs. The $t$-test is performed on paired inputs of the subnetwork performance and ESM-2 baseline performance and $p$-values are reported below. The random residue control subnetwork is trained to suppress random residues, but we independently evaluate and report the MLM performance of this subnetwork on alpha helices and beta sheets. All per-sequence metrics are available in CSVs in the code repository.

| Category | Target | Sparsity | Training Step | Subnet. Supp. | Subnet. Maint. | ESM-2 Supp. | ESM-2 Maint. | t-Test ($p$) Supp. | t-Test ($p$) Maint. |
|---|---|---|---|---|---|---|---|---|---|
| Residue | Helix | 0.90 | 784 | $7.9 \pm 5.0$ | $5.0 \pm 5.2$ | $5.6 \pm 4.0$ | $4.9 \pm 5.1$ | $< 1e{-16}$ | $< 1e{-16}$ |
| | Sheet | 0.89 | 784 | $6.7 \pm 6.1$ | $5.6 \pm 4.1$ | $4.9 \pm 5.1$ | $5.6 \pm 4.0$ | $< 1e{-16}$ | $< 1e{-16}$ |
| Class | 1 | 2.13 | 7,680 | $27.2 \pm 80.3$ | $8.0 \pm 15.4$ | $8.6 \pm 15.2$ | $8.8 \pm 14.1$ | $< 1e{-16}$ | $< 1e{-16}$ |
| | 2 | 2.06 | 7,200 | $28.7 \pm 74.1$ | $8.2 \pm 15.5$ | $8.9 \pm 15.7$ | $8.7 \pm 14.0$ | $< 1e{-16}$ | $< 1e{-16}$ |

| Category | Target | Sparsity | Training Step | Subnet. Supp. | Subnet. Maint. | ESM-2 Supp. | ESM-2 Maint. | t-Test ($p$) Supp. | t-Test ($p$) Maint. |
|---|---|---|---|---|---|---|---|---|---|
| | 3 | 2.53 | 7,820 | 41.5±91.4 | 8.2 ± 18.5 | 8.9 ± 13.7 | 8.6 ± 15.2 | $< 1e{-16}$ | $< 1e{-16}$ |
| Architecture | 1.10 | 2.02 | 4,440 | 11.0±21.7 | 9.3 ± 16.4 | 7.6 ± 15.2 | 8.9 ± 14.3 | $< 1e{-16}$ | $< 1e{-16}$ |
| | 1.20 | 1.25 | 3,660 | 14.1±28.0 | 9.3 ± 17.6 | 9.0 ± 12.4 | 8.7 ± 14.5 | $< 1e{-16}$ | $< 1e{-16}$ |
| | 1.25 | 1.31 | 640 | 34.9±70.6 | 9.1 ± 16.8 | 11.9±19.6 | 8.7 ± 14.2 | $< 1e{-16}$ | $< 1e{-16}$ |
| | 2.30 | 1.05 | 1,680 | 9.9 ± 12.7 | 9.5 ± 17.0 | 5.1 ± 5.1 | 8.8 ± 14.5 | $< 1e{-16}$ | $< 1e{-16}$ |
| | 2.40 | 1.10 | 2,280 | 13.0±21.4 | 9.3 ± 17.1 | 7.2 ± 10.5 | 8.8 ± 14.6 | $< 1e{-16}$ | $< 1e{-16}$ |
| | 2.60 | 1.71 | 3,000 | 20.0±37.6 | 9.2 ± 16.5 | 8.9 ± 14.1 | 8.7 ± 14.4 | $< 1e{-16}$ | $< 1e{-16}$ |
| | 3.30 | 1.68 | 4,880 | 10.1±14.6 | 9.3 ± 17.6 | 7.3 ± 10.6 | 9.0 ± 15.0 | $< 1e{-16}$ | $< 1e{-16}$ |
| | 3.40 | 2.18 | 6,200 | 34.9±67.5 | 8.8 ± 18.2 | 9.2 ± 14.0 | 8.6 ± 14.5 | $< 1e{-16}$ | $< 1e{-16}$ |
| | 3.90 | 1.41 | 1,640 | 14.9±28.0 | 9.0 ± 16.3 | 10.8±19.4 | 8.6 ± 14.1 | $< 1e{-16}$ | $< 1e{-16}$ |
| Topology | 1.10.10 | 1.51 | 1,280 | 6.9 ± 9.6 | 9.7 ± 17.5 | 4.6 ± 6.7 | 8.8 ± 14.5 | $< 1e{-16}$ | $< 1e{-16}$ |
| | 1.10.287 | 1.37 | 720 | 5.3 ± 5.7 | 9.4 ± 17.1 | 5.2 ± 6.1 | 8.8 ± 14.5 | $4.5e-01$ | $< 1e{-16}$ |
| | 1.20.120 | 1.38 | 800 | 11.9±14.5 | 9.5 ± 16.9 | 8.3 ± 8.5 | 8.7 ± 14.5 | $< 1e{-16}$ | $< 1e{-16}$ |
| | 1.20.58 | 1.41 | 960 | 13.4±19.8 | 9.6 ± 17.3 | 8.4 ± 10.8 | 8.7 ± 14.4 | $< 1e{-16}$ | $< 1e{-16}$ |
| | 1.25.40 | 1.34 | 780 | 40.8±94.2 | 9.2 ± 16.7 | 11.0±19.2 | 8.7 ± 14.3 | $< 1e{-16}$ | $< 1e{-16}$ |
| | 2.40.50 | 1.36 | 720 | 8.7 ± 9.7 | 9.3 ± 16.4 | 5.2 ± 5.8 | 8.8 ± 14.5 | $< 1e{-16}$ | $< 1e{-16}$ |
| | 2.60.120 | 1.37 | 1,000 | 32.7±49.6 | 9.1 ± 17.9 | 11.1±15.0 | 8.7 ± 14.4 | $< 1e{-16}$ | $< 1e{-16}$ |
| | 2.60.40 | 1.52 | 1,800 | 11.9±15.3 | 9.4 ± 17.5 | 7.1 ± 9.7 | 8.8 ± 14.6 | $< 1e{-16}$ | $< 1e{-16}$ |
| | 3.30.70 | 1.52 | 1,500 | 8.3 ± 9.4 | 9.6 ± 18.1 | 5.7 ± 6.3 | 8.8 ± 14.6 | $< 1e{-16}$ | $< 1e{-16}$ |
| | 3.40.50 | 1.80 | 3,840 | 25.6±40.2 | 9.1 ± 17.7 | 8.8 ± 11.1 | 8.7 ± 14.8 | $< 1e{-16}$ | $< 1e{-16}$ |
| H. Superfam. | 1.10.10.10 | 1.26 | 300 | 4.0 ± 3.4 | 9.0 ± 16.5 | 4.1 ± 3.9 | 8.8 ± 14.5 | $4.1e-02$ | $< 1e{-16}$ |
| | 2.40.50.140 | 1.38 | 480 | 10.2±11.0 | 9.5 ± 17.4 | 4.6 ± 3.6 | 8.8 ± 14.4 | $< 1e{-16}$ | $< 1e{-16}$ |
| | 2.60.40.10 | 1.04 | 840 | 8.8 ± 9.1 | 9.2 ± 16.4 | 4.9 ± 4.3 | 8.8 ± 14.5 | $< 1e{-16}$ | $< 1e{-16}$ |
| | 3.20.20.80 | 1.26 | 360 | 46.4±55.6 | 9.1 ± 16.8 | 12.9±14.5 | 8.7 ± 14.4 | $< 1e{-16}$ | $< 1e{-16}$ |
| | 3.40.190.10 | 1.29 | 420 | 22.7±30.1 | 9.2 ± 16.5 | 8.9 ± 8.6 | 8.7 ± 14.4 | $< 1e{-16}$ | $< 1e{-16}$ |
| | 3.40.30.10 | 1.32 | 480 | 15.1±13.5 | 9.3 ± 17.5 | 5.1 ± 4.4 | 8.8 ± 14.4 | $< 1e{-16}$ | $< 1e{-16}$ |
| | 3.40.50.150 | 1.31 | 480 | 17.5±24.0 | 9.2 ± 17.5 | 9.5 ± 12.1 | 8.7 ± 14.4 | $< 1e{-16}$ | $< 1e{-16}$ |
| | 3.40.50.1820 | 1.27 | 360 | 20.2±21.8 | 9.1 ± 16.5 | 12.2±12.6 | 8.7 ± 14.4 | $< 1e{-16}$ | $< 1e{-16}$ |
| | 3.40.50.300 | 1.30 | 560 | 21.6±28.1 | 9.1 ± 16.3 | 9.2 ± 10.2 | 8.7 ± 14.5 | $< 1e{-16}$ | $< 1e{-16}$ |
| | 3.40.50.720 | 1.32 | 660 | 27.6±49.2 | 9.2 ± 16.7 | 7.4 ± 7.8 | 8.8 ± 14.5 | $< 1e{-16}$ | $< 1e{-16}$ |
| Random | Helix | 0.87 | 196 | 6.2 ± 4.3 | 5.2 ± 5.2 | 5.6 ± 4.0 | 4.9 ± 5.1 | $< 1e{-16}$ | $< 1e{-16}$ |
| | Sheet | 0.87 | 196 | 5.2 ± 5.2 | 6.2 ± 4.3 | 4.9 ± 5.1 | 5.6 ± 4.0 | $< 1e{-16}$ | $< 1e{-16}$ |
| | 100 seqs. | 1.17 | 420 | 9.2 ± 13.2 | 8.9 ± 15.7 | 8.4 ± 12.0 | 8.7 ± 14.4 | $1.8e-09$ | $< 1e{-16}$ |
| | 200 seqs. | 1.21 | 780 | 9.2 ± 13.2 | 9.0 ± 15.7 | 8.3 ± 11.8 | 8.7 ± 14.4 | $< 1e{-16}$ | $< 1e{-16}$ |
| | 1000 seqs. | 1.37 | 2,520 | 8.5 ± 13.4 | 8.9 ± 16.2 | 7.8 ± 11.5 | 8.8 ± 14.6 | $< 1e{-16}$ | $2.1e-04$ |
| | 2000 seqs. | 2.17 | 7,500 | 11.2±21.3 | 8.2 ± 15.6 | 8.6 ± 14.0 | 8.8 ± 14.5 | $< 1e{-16}$ | $< 1e{-16}$ |

**Table S5. ProteinMPNN perplexities on CATH PDBs (Mainly Alpha, Mainly Beta, and Alpha-Beta classes).** Reported values correspond to mean, standard deviation, minimum, and maximum perplexities across all CATH domains within each structural CATH Class.

| Category | Count (n) | Mean | Std. Dev. | Min | Max |
|---|---|---|---|---|---|
| Mainly Alpha | 10506 | 4.49 | 3.02 | 1.51 | 44.00 |
| Mainly Beta | 8493 | 3.71 | 2.15 | 1.56 | 35.19 |
| Alpha-Beta | 19092 | 3.50 | 1.97 | 1.65 | 57.61 |

**Table S6. ESM-2 650M subnetwork and baseline structure prediction performance using the ESMFold (650M) folding trunk on the validation datasets**. For TM-score, RMSD, and pLDDT, we report the mean ± standard deviation of the subnetwork performance across all sequences within each category of suppression and maintenance inputs. ESM-2 (650M) performance on the same categories is reported as the PLM baseline. To quantify the differences in subnetwork and PLM performance, we perform a paired t-test on all (i) suppression inputs and (ii) maintenance inputs, computing the difference in $\Delta\text{metric} = \text{metric}_{\text{Subnet.}} - \text{metric}_{\text{ESM-2.}}$. We then perform a Kolmogorov–Smirnov (KS) test on these differences to assess whether the distribution of $|\Delta_{\text{metric,supp}}|$ is significantly greater than that of $|\Delta_{\text{metric,maint}}|$. Our evaluation scheme is illustrated in Fig. 3A. We report $p$-values for both statistical tests for each subnetwork; significant $p$-values are in bold. For residue-level suppression, we evaluate structure prediction on CATH class categories of mainly alpha and mainly beta sequences as a proxy for evaluating alpha helix and beta strand performance. We report performance on the only residue-specific metric of pLDDT in Fig. 3F-G. The random residue suppression subnetwork, i.e. residue-control, is one subnetwork but we evaluated it separately on alpha helices and beta sheets. All per-sequence structure prediction metrics are provided via CSVs in our code repository.

| Category | Target | Metric | Subnet. Supp. | Subnet. Maint. | ESM-2 Supp. | ESM-2 Maint. | t-Test ($p$) Supp. | t-Test ($p$) Maint. | K.S.-test ($p$) |
|---|---|---|---|---|---|---|---|---|---|
| Residue | Helix | RMSD | 3.62 ± 1.19 | 3.05 ± 1.09 | 2.93 ± 1.07 | 2.83 ± 1.09 | **< 1e−16** | **< 1e−16** | **< 1e−16** |
| | | TM-score | 0.55 ± 0.20 | 0.68 ± 0.19 | 0.66 ± 0.20 | 0.71 ± 0.19 | **< 1e−16** | **< 1e−16** | **< 1e−16** |
| | | pLDDT | 50.81±13.89 | 59.49±12.79 | 66.51±12.82 | 65.38±12.69 | **< 1e−16** | **< 1e−16** | **< 1e−16** |
| | Sheet | RMSD | 3.74 ± 1.43 | 3.05 ± 1.12 | 2.89 ± 1.19 | 2.85 ± 1.11 | **< 1e−16** | **< 1e−16** | **< 1e−16** |
| | | TM-score | 0.57 ± 0.21 | 0.66 ± 0.20 | 0.70 ± 0.19 | 0.69 ± 0.20 | **< 1e−16** | **< 1e−16** | **< 1e−16** |
| | | pLDDT | 47.79±13.55 | 60.98±12.72 | 63.86±13.49 | 65.87±12.65 | **< 1e−16** | **< 1e−16** | **< 1e−16** |
| Class | 1 | RMSD | 5.22 ± 1.42 | 3.05 ± 1.10 | 2.86 ± 1.12 | 2.82 ± 1.10 | **< 1e−16** | **< 1e−16** | **< 1e−16** |
| | | TM-score | 0.27 ± 0.12 | 0.68 ± 0.19 | 0.66 ± 0.21 | 0.71 ± 0.20 | **< 1e−16** | **< 1e−16** | **< 1e−16** |
| | | pLDDT | 33.31 ± 8.37 | 62.30±13.07 | 67.09±13.15 | 65.24±12.92 | **< 1e−16** | **< 1e−16** | **< 1e−16** |
| | 2 | RMSD | 6.02 ± 1.28 | 3.08 ± 1.10 | 2.93 ± 1.15 | 2.84 ± 1.09 | **< 1e−16** | **< 1e−16** | **< 1e−16** |
| | | TM-score | 0.23 ± 0.11 | 0.66 ± 0.20 | 0.69 ± 0.19 | 0.69 ± 0.20 | **< 1e−16** | **< 1e−16** | **< 1e−16** |
| | | pLDDT | 30.53 ± 7.66 | 64.25±12.91 | 63.02±13.59 | 66.16±12.66 | **< 1e−16** | **< 1e−16** | **< 1e−16** |
| | 3 | RMSD | 6.13 ± 1.39 | 3.26 ± 1.15 | 2.81 ± 1.06 | 2.93 ± 1.13 | **< 1e−16** | **< 1e−16** | **< 1e−16** |
| | | TM-score | 0.23 ± 0.12 | 0.61 ± 0.20 | 0.72 ± 0.19 | 0.67 ± 0.21 | **< 1e−16** | **< 1e−16** | **< 1e−16** |
| | | pLDDT | 30.80 ± 7.97 | 60.54±13.99 | 65.64±12.55 | 64.75±13.30 | **< 1e−16** | **< 1e−16** | **< 1e−16** |
| Arch. | 1.10 | RMSD | 4.80 ± 1.20 | 3.12 ± 1.16 | 2.85 ± 1.13 | 2.83 ± 1.11 | **< 1e−16** | **< 1e−16** | **< 1e−16** |
| | | TM-score | 0.32 ± 0.13 | 0.66 ± 0.21 | 0.65 ± 0.20 | 0.71 ± 0.20 | **< 1e−16** | **< 1e−16** | **< 1e−16** |
| | | pLDDT | 35.91 ± 8.20 | 61.82±14.58 | 66.94±12.45 | 65.53±12.96 | **< 1e−16** | **< 1e−16** | **< 1e−16** |
| | 1.20 | RMSD | 4.59 ± 1.27 | 3.05 ± 1.16 | 3.06 ± 1.08 | 2.84 ± 1.10 | **< 1e−16** | **< 1e−16** | **< 1e−16** |
| | | TM-score | 0.36 ± 0.17 | 0.66 ± 0.21 | 0.63 ± 0.20 | 0.70 ± 0.20 | **< 1e−16** | **< 1e−16** | **< 1e−16** |
| | | pLDDT | 39.83±13.38 | 62.42±14.80 | 65.69±13.68 | 65.36±13.01 | **< 1e−16** | **< 1e−16** | **< 1e−16** |
| | 1.25 | RMSD | 4.29 ± 1.40 | 3.00 ± 1.10 | 2.67 ± 1.04 | 2.86 ± 1.10 | **4.6e-12** | **< 1e−16** | **< 1e−16** |

| Category | Target | Metric | Subnet. Supp. | Subnet. Maint. | ESM-2 Supp. | ESM-2 Maint. | t-Test ($p$) Supp. | t-Test ($p$) Maint. | K.S.-test ($p$) |
|---|---|---|---|---|---|---|---|---|---|
| | | TM-score | $0.54 \pm 0.22$ | $0.67 \pm 0.20$ | $0.78 \pm 0.15$ | $0.69 \pm 0.20$ | **7.5e-12** | **< 1e−16** | **< 1e−16** |
| | | pLDDT | $49.23 \pm 15.56$ | $63.32 \pm 13.03$ | $71.39 \pm 11.65$ | $65.30 \pm 12.65$ | **5.7e-16** | **< 1e−16** | **< 1e−16** |
| | 2.30 | RMSD | $4.66 \pm 1.21$ | $3.06 \pm 1.17$ | $2.69 \pm 0.95$ | $2.86 \pm 1.12$ | **< 1e−16** | **< 1e−16** | **< 1e−16** |
| | | TM-score | $0.37 \pm 0.18$ | $0.67 \pm 0.20$ | $0.68 \pm 0.18$ | $0.70 \pm 0.20$ | **< 1e−16** | **< 1e−16** | **< 1e−16** |
| | | pLDDT | $37.72 \pm 12.77$ | $62.99 \pm 14.17$ | $67.14 \pm 13.07$ | $65.61 \pm 13.01$ | **< 1e−16** | **< 1e−16** | **< 1e−16** |
| | 2.40 | RMSD | $5.12 \pm 1.45$ | $3.00 \pm 1.15$ | $2.99 \pm 1.07$ | $2.81 \pm 1.09$ | **< 1e−16** | **< 1e−16** | **< 1e−16** |
| | | TM-score | $0.34 \pm 0.20$ | $0.67 \pm 0.20$ | $0.66 \pm 0.19$ | $0.70 \pm 0.20$ | **< 1e−16** | **< 1e−16** | **< 1e−16** |
| | | pLDDT | $35.83 \pm 12.39$ | $63.38 \pm 13.86$ | $62.20 \pm 14.02$ | $66.23 \pm 12.59$ | **< 1e−16** | **< 1e−16** | **< 1e−16** |
| | 2.60 | RMSD | $6.03 \pm 1.09$ | $3.07 \pm 1.15$ | $2.84 \pm 1.16$ | $2.86 \pm 1.11$ | **< 1e−16** | **< 1e−16** | **< 1e−16** |
| | | TM-score | $0.24 \pm 0.12$ | $0.66 \pm 0.21$ | $0.71 \pm 0.18$ | $0.69 \pm 0.20$ | **< 1e−16** | **< 1e−16** | **< 1e−16** |
| | | pLDDT | $29.96 \pm 6.60$ | $63.17 \pm 13.73$ | $63.26 \pm 13.72$ | $65.66 \pm 12.94$ | **< 1e−16** | **< 1e−16** | **< 1e−16** |
| | 3.30 | RMSD | $4.93 \pm 1.24$ | $3.05 \pm 1.14$ | $2.87 \pm 1.14$ | $2.80 \pm 1.09$ | **< 1e−16** | **< 1e−16** | **< 1e−16** |
| | | TM-score | $0.34 \pm 0.16$ | $0.67 \pm 0.20$ | $0.66 \pm 0.20$ | $0.71 \pm 0.20$ | **< 1e−16** | **< 1e−16** | **< 1e−16** |
| | | pLDDT | $36.39 \pm 11.38$ | $63.25 \pm 13.87$ | $64.26 \pm 14.23$ | $66.24 \pm 12.61$ | **< 1e−16** | **< 1e−16** | **< 1e−16** |
| | 3.40 | RMSD | $6.42 \pm 1.24$ | $3.18 \pm 1.14$ | $2.60 \pm 0.98$ | $2.92 \pm 1.13$ | **< 1e−16** | **< 1e−16** | **< 1e−16** |
| | | TM-score | $0.24 \pm 0.11$ | $0.63 \pm 0.20$ | $0.77 \pm 0.16$ | $0.67 \pm 0.20$ | **< 1e−16** | **< 1e−16** | **< 1e−16** |
| | | pLDDT | $30.33 \pm 6.87$ | $61.31 \pm 13.65$ | $68.47 \pm 10.68$ | $64.63 \pm 13.30$ | **< 1e−16** | **< 1e−16** | **< 1e−16** |
| | 3.90 | RMSD | $4.85 \pm 1.49$ | $3.10 \pm 1.22$ | $3.32 \pm 1.38$ | $2.84 \pm 1.11$ | **< 1e−16** | **< 1e−16** | **< 1e−16** |
| | | TM-score | $0.42 \pm 0.21$ | $0.66 \pm 0.21$ | $0.65 \pm 0.23$ | $0.70 \pm 0.20$ | **< 1e−16** | **< 1e−16** | **< 1e−16** |
| | | pLDDT | $41.17 \pm 12.83$ | $61.81 \pm 15.13$ | $61.12 \pm 13.68$ | $65.50 \pm 13.06$ | **< 1e−16** | **< 1e−16** | **< 1e−16** |
| Topo. | 1.10.10 | RMSD | $3.90 \pm 1.10$ | $3.05 \pm 1.15$ | $2.35 \pm 0.90$ | $2.87 \pm 1.13$ | **9.0e-14** | **< 1e−16** | **< 1e−16** |
| | | TM-score | $0.41 \pm 0.16$ | $0.67 \pm 0.20$ | $0.68 \pm 0.18$ | $0.70 \pm 0.20$ | **1.3e-14** | **< 1e−16** | **< 1e−16** |
| | | pLDDT | $44.81 \pm 14.34$ | $62.72 \pm 14.01$ | $71.96 \pm 10.84$ | $65.30 \pm 13.04$ | **< 1e−16** | **< 1e−16** | **< 1e−16** |
| | 1.10.287 | RMSD | $2.80 \pm 0.93$ | $2.98 \pm 1.08$ | $2.77 \pm 0.86$ | $2.85 \pm 1.08$ | 7.6e-01 | **< 1e−16** | **3.2e-02** |
| | | TM-score | $0.56 \pm 0.17$ | $0.68 \pm 0.19$ | $0.59 \pm 0.18$ | $0.70 \pm 0.20$ | **6.2e-03** | **< 1e−16** | 6.1e-02 |
| | | pLDDT | $67.32 \pm 13.60$ | $63.70 \pm 12.75$ | $70.62 \pm 11.32$ | $65.74 \pm 12.57$ | **5.9e-03** | **< 1e−16** | **4.6e-02** |
| | 1.20.120 | RMSD | $4.10 \pm 1.35$ | $3.00 \pm 1.11$ | $3.28 \pm 1.13$ | $2.83 \pm 1.09$ | **1.4e-04** | **< 1e−16** | **2.8e-05** |
| | | TM-score | $0.47 \pm 0.19$ | $0.67 \pm 0.20$ | $0.61 \pm 0.20$ | $0.70 \pm 0.20$ | **7.8e-07** | **< 1e−16** | **9.6e-11** |
| | | pLDDT | $48.32 \pm 14.04$ | $63.08 \pm 13.69$ | $63.44 \pm 12.66$ | $65.64 \pm 12.96$ | **1.6e-07** | **< 1e−16** | **1.2e-10** |
| | 1.20.58 | RMSD | $3.40 \pm 0.98$ | $3.00 \pm 1.10$ | $2.93 \pm 1.13$ | $2.85 \pm 1.10$ | **3.3e-04** | **< 1e−16** | **4.3e-04** |
| | | TM-score | $0.50 \pm 0.16$ | $0.68 \pm 0.20$ | $0.60 \pm 0.21$ | $0.70 \pm 0.20$ | **1.4e-05** | **< 1e−16** | **5.6e-08** |
| | | pLDDT | $55.58 \pm 12.59$ | $63.37 \pm 13.01$ | $64.98 \pm 14.31$ | $65.58 \pm 12.65$ | **7.5e-06** | **< 1e−16** | **5.6e-07** |
| | 1.25.40 | RMSD | $4.21 \pm 1.60$ | $2.98 \pm 1.13$ | $2.65 \pm 1.14$ | $2.85 \pm 1.11$ | **1.3e-08** | **< 1e−16** | **1.0e-12** |
| | | TM-score | $0.49 \pm 0.25$ | $0.68 \pm 0.20$ | $0.77 \pm 0.15$ | $0.70 \pm 0.20$ | **5.2e-10** | **< 1e−16** | **2.1e-16** |
| | | pLDDT | $47.61 \pm 17.00$ | $63.50 \pm 13.65$ | $74.72 \pm 10.08$ | $65.48 \pm 13.00$ | **1.7e-13** | **< 1e−16** | **< 1e−16** |
| | 2.40.50 | RMSD | $4.66 \pm 1.19$ | $2.98 \pm 1.12$ | $2.86 \pm 1.11$ | $2.86 \pm 1.10$ | **1.1e-11** | **< 1e−16** | **< 1e−16** |
| | | TM-score | $0.35 \pm 0.17$ | $0.68 \pm 0.20$ | $0.63 \pm 0.21$ | $0.69 \pm 0.20$ | **6.2e-11** | **< 1e−16** | **< 1e−16** |
| | | pLDDT | $37.35 \pm 10.71$ | $63.53 \pm 13.34$ | $62.32 \pm 14.64$ | $65.39 \pm 12.84$ | **2.3e-12** | **< 1e−16** | **< 1e−16** |
| | 2.60.120 | RMSD | $5.75 \pm 1.65$ | $3.00 \pm 1.15$ | $2.91 \pm 1.15$ | $2.86 \pm 1.12$ | **< 1e−16** | **< 1e−16** | **< 1e−16** |
| | | TM-score | $0.36 \pm 0.22$ | $0.67 \pm 0.21$ | $0.74 \pm 0.17$ | $0.69 \pm 0.20$ | **< 1e−16** | **< 1e−16** | **< 1e−16** |
| | | pLDDT | $33.70 \pm 10.54$ | $63.74 \pm 13.81$ | $62.51 \pm 12.81$ | $65.47 \pm 13.04$ | **< 1e−16** | **< 1e−16** | **< 1e−16** |
| | 2.60.40 | RMSD | $5.29 \pm 1.24$ | $3.02 \pm 1.14$ | $2.61 \pm 0.91$ | $2.85 \pm 1.10$ | **< 1e−16** | **< 1e−16** | **< 1e−16** |
| | | TM-score | $0.32 \pm 0.17$ | $0.67 \pm 0.20$ | $0.73 \pm 0.15$ | $0.69 \pm 0.20$ | **< 1e−16** | **< 1e−16** | **< 1e−16** |
| | | pLDDT | $32.83 \pm 8.78$ | $64.14 \pm 13.58$ | $65.96 \pm 12.34$ | $65.44 \pm 12.82$ | **< 1e−16** | **< 1e−16** | **< 1e−16** |
| | 3.30.70 | RMSD | $4.12 \pm 1.11$ | $3.05 \pm 1.14$ | $2.72 \pm 1.13$ | $2.84 \pm 1.11$ | **< 1e−16** | **< 1e−16** | **< 1e−16** |
| | | TM-score | $0.44 \pm 0.18$ | $0.67 \pm 0.20$ | $0.67 \pm 0.20$ | $0.70 \pm 0.20$ | **< 1e−16** | **< 1e−16** | **< 1e−16** |
| | | pLDDT | $41.78 \pm 13.24$ | $62.50 \pm 13.90$ | $65.82 \pm 14.80$ | $65.76 \pm 12.80$ | **< 1e−16** | **< 1e−16** | **< 1e−16** |
| | 3.40.50 | RMSD | $5.99 \pm 1.16$ | $3.09 \pm 1.13$ | $2.53 \pm 0.88$ | $2.91 \pm 1.14$ | **< 1e−16** | **< 1e−16** | **< 1e−16** |
| | | TM-score | $0.32 \pm 0.14$ | $0.66 \pm 0.20$ | $0.80 \pm 0.14$ | $0.68 \pm 0.20$ | **< 1e−16** | **< 1e−16** | **< 1e−16** |
| | | pLDDT | $32.65 \pm 6.56$ | $62.69 \pm 13.36$ | $69.77 \pm 9.16$ | $64.73 \pm 13.38$ | **< 1e−16** | **< 1e−16** | **< 1e−16** |
| H. Supfam. | 1.10.10.10 | RMSD | $2.30 \pm 0.99$ | $2.92 \pm 1.10$ | $2.28 \pm 0.89$ | $2.83 \pm 1.09$ | 8.8e-01 | **< 1e−16** | 1.8e-01 |
| | | TM-score | $0.71 \pm 0.17$ | $0.69 \pm 0.20$ | $0.72 \pm 0.16$ | $0.70 \pm 0.20$ | 4.7e-01 | **< 1e−16** | 7.0e-02 |
| | | pLDDT | $72.62 \pm 11.95$ | $65.18 \pm 12.92$ | $73.16 \pm 11.74$ | $65.74 \pm 12.81$ | 9.3e-02 | **< 1e−16** | 4.5e-01 |
| | 2.40.50.140 | RMSD | $5.03 \pm 0.96$ | $2.99 \pm 1.09$ | $2.49 \pm 0.90$ | $2.85 \pm 1.08$ | **1.1e-08** | **< 1e−16** | **< 1e−16** |
| | | TM-score | $0.32 \pm 0.14$ | $0.68 \pm 0.20$ | $0.72 \pm 0.17$ | $0.70 \pm 0.20$ | **6.3e-08** | **< 1e−16** | **2.4e-14** |

| Category | Target | Metric | Subnet. Supp. | Subnet. Maint. | ESM-2 Supp. | ESM-2 Maint. | t-Test ($p$) Supp. | t-Test ($p$) Maint. | K.S.-test ($p$) |
|---|---|---|---|---|---|---|---|---|---|
| | | pLDDT | 34.61 ± 8.13 | 63.92±13.16 | 68.78±12.37 | 65.73±12.64 | **1.1e-07** | **< 1e−16** | **< 1e−16** |
| | 2.60.40.10 | RMSD | 3.75 ± 1.26 | 3.03 ± 1.14 | 2.17 ± 0.69 | 2.89 ± 1.14 | **7.2e-13** | **< 1e−16** | **< 1e−16** |
| | | TM-score | 0.53 ± 0.19 | 0.67 ± 0.20 | 0.79 ± 0.10 | 0.69 ± 0.20 | **5.4e-15** | **< 1e−16** | **< 1e−16** |
| | | pLDDT | 41.57±12.07 | 63.52±13.34 | 71.28 ± 9.64 | 65.13±13.02 | **< 1e−16** | **< 1e−16** | **< 1e−16** |
| | 3.20.20.80 | RMSD | 3.45 ± 0.94 | 2.96 ± 1.11 | 2.63 ± 0.66 | 2.86 ± 1.10 | **6.4e-06** | **< 1e−16** | **1.8e-07** |
| | | TM-score | 0.79 ± 0.10 | 0.68 ± 0.20 | 0.87 ± 0.07 | 0.69 ± 0.20 | **1.2e-05** | **< 1e−16** | **6.3e-08** |
| | | pLDDT | 49.76 ± 9.16 | 63.98±13.30 | 67.03 ± 7.20 | 65.46±12.91 | **8.5e-10** | **< 1e−16** | **< 1e−16** |
| | 3.40.190.10 | RMSD | 5.22 ± 1.21 | 3.00 ± 1.14 | 2.84 ± 1.06 | 2.89 ± 1.15 | **5.1e-11** | **< 1e−16** | **< 1e−16** |
| | | TM-score | 0.32 ± 0.11 | 0.67 ± 0.20 | 0.63 ± 0.12 | 0.69 ± 0.20 | **2.6e-09** | **< 1e−16** | **< 1e−16** |
| | | pLDDT | 32.96 ± 5.12 | 64.06±13.44 | 66.66 ± 6.61 | 65.12±13.23 | **9.4e-16** | **< 1e−16** | **< 1e−16** |
| | 3.40.30.10 | RMSD | 4.49 ± 1.63 | 2.95 ± 1.09 | 1.75 ± 0.48 | 2.86 ± 1.11 | **1.7e-06** | **< 1e−16** | **1.2e-14** |
| | | TM-score | 0.46 ± 0.21 | 0.68 ± 0.20 | 0.84 ± 0.08 | 0.69 ± 0.20 | **1.7e-06** | **< 1e−16** | **2.5e-15** |
| | | pLDDT | 41.38±14.66 | 64.21±12.76 | 76.66 ± 7.22 | 65.51±12.75 | **1.7e-09** | **< 1e−16** | **< 1e−16** |
| | 3.40.50.150 | RMSD | 2.85 ± 1.23 | 2.94 ± 1.11 | 2.40 ± 0.77 | 2.84 ± 1.10 | **2.2e-03** | **< 1e−16** | **1.5e-03** |
| | | TM-score | 0.79 ± 0.16 | 0.68 ± 0.20 | 0.85 ± 0.10 | 0.70 ± 0.20 | **5.8e-04** | **< 1e−16** | **2.8e-04** |
| | | pLDDT | 62.15±12.19 | 63.90±12.84 | 72.03 ± 9.41 | 65.70±12.82 | **8.7e-10** | **< 1e−16** | **< 1e−16** |
| | 3.40.50.1820 | RMSD | 2.57 ± 0.70 | 2.95 ± 1.13 | 2.39 ± 0.68 | 2.87 ± 1.12 | **4.9e-05** | **< 1e−16** | **1.7e-02** |
| | | TM-score | 0.86 ± 0.07 | 0.68 ± 0.20 | 0.88 ± 0.06 | 0.69 ± 0.20 | **2.9e-06** | **< 1e−16** | **2.5e-03** |
| | | pLDDT | 66.94 ± 8.82 | 64.39±13.30 | 71.50 ± 6.95 | 65.29±13.09 | **7.4e-08** | **< 1e−16** | **8.1e-10** |
| | 3.40.50.300 | RMSD | 3.44 ± 1.11 | 2.99 ± 1.12 | 2.63 ± 0.76 | 2.87 ± 1.12 | **2.0e-08** | **< 1e−16** | **1.7e-10** |
| | | TM-score | 0.68 ± 0.15 | 0.67 ± 0.20 | 0.79 ± 0.13 | 0.69 ± 0.20 | **2.2e-08** | **< 1e−16** | **3.9e-12** |
| | | pLDDT | 50.62±12.41 | 63.22±13.53 | 68.34 ± 8.00 | 65.36±13.26 | **< 1e−16** | **< 1e−16** | **< 1e−16** |
| | 3.40.50.720 | RMSD | 4.26 ± 1.41 | 2.98 ± 1.11 | 2.47 ± 0.93 | 2.87 ± 1.11 | **3.5e-08** | **< 1e−16** | **< 1e−16** |
| | | TM-score | 0.56 ± 0.19 | 0.68 ± 0.20 | 0.77 ± 0.16 | 0.70 ± 0.20 | **5.4e-09** | **< 1e−16** | **< 1e−16** |
| | | pLDDT | 42.35±11.23 | 65.14±13.51 | 72.20±10.85 | 65.42±12.87 | **1.2e-14** | **1.3e-04** | **< 1e−16** |
| Random | 100 seqs. | RMSD | 3.06 ± 1.17 | 2.92 ± 1.11 | 2.92 ± 1.17 | 2.84 ± 1.11 | **1.0e-03** | **< 1e−16** | 5.2e-01 |
| | | TM-score | 0.66 ± 0.20 | 0.69 ± 0.20 | 0.67 ± 0.21 | 0.70 ± 0.20 | **3.7e-04** | **< 1e−16** | 5.8e-01 |
| | | pLDDT | 63.15±12.98 | 65.23±13.21 | 64.74±12.34 | 65.71±12.95 | **3.4e-07** | **< 1e−16** | **5.3e-06** |
| | 1000 seqs. | RMSD | 3.64 ± 1.24 | 3.37 ± 1.25 | 2.81 ± 1.05 | 2.86 ± 1.12 | **< 1e−16** | **< 1e−16** | **< 1e−16** |
| | | TM-score | 0.57 ± 0.21 | 0.62 ± 0.21 | 0.70 ± 0.19 | 0.69 ± 0.20 | **< 1e−16** | **< 1e−16** | **< 1e−16** |
| | | pLDDT | 49.65±13.98 | 55.49±14.77 | 65.79±12.85 | 65.32±13.06 | **< 1e−16** | **< 1e−16** | **< 1e−16** |
| | 200 seqs. | RMSD | 3.00 ± 1.16 | 2.95 ± 1.13 | 2.79 ± 1.12 | 2.85 ± 1.13 | **4.6e-10** | **< 1e−16** | **1.5e-03** |
| | | TM-score | 0.67 ± 0.20 | 0.68 ± 0.20 | 0.70 ± 0.20 | 0.70 ± 0.20 | **2.4e-12** | **< 1e−16** | **1.6e-04** |
| | | pLDDT | 63.04±14.38 | 64.62±13.40 | 65.62±13.21 | 65.50±13.01 | **5.6e-13** | **< 1e−16** | **1.3e-06** |
| | 2000 seqs. | RMSD | 5.21 ± 1.30 | 3.93 ± 1.48 | 2.85 ± 1.08 | 2.86 ± 1.12 | **< 1e−16** | **< 1e−16** | **< 1e−16** |
| | | TM-score | 0.34 ± 0.15 | 0.53 ± 0.22 | 0.70 ± 0.20 | 0.69 ± 0.20 | **< 1e−16** | **< 1e−16** | **< 1e−16** |
| | | pLDDT | 34.59 ± 6.67 | 48.86±15.22 | 65.59±12.74 | 65.41±13.08 | **< 1e−16** | **< 1e−16** | **< 1e−16** |
| | Helix | RMSD | 3.23 ± 1.17 | 3.02 ± 1.12 | 2.92 ± 1.14 | 2.81 ± 1.09 | **< 1e−16** | **< 1e−16** | **2.1e-08** |
| | | TM-score | 0.60 ± 0.20 | 0.68 ± 0.20 | 0.65 ± 0.21 | 0.71 ± 0.19 | **< 1e−16** | **< 1e−16** | **1.4e-12** |
| | | pLDDT | 59.77±14.46 | 60.31±13.52 | 66.43±13.05 | 65.37±12.75 | **< 1e−16** | **< 1e−16** | **1.6e-13** |
| | Sheet | RMSD | 3.04 ± 1.20 | 3.08 ± 1.12 | 2.89 ± 1.17 | 2.82 ± 1.08 | **1.0e-16** | **< 1e−16** | **4.9e-09** |
| | | TM-score | 0.67 ± 0.20 | 0.66 ± 0.20 | 0.70 ± 0.19 | 0.70 ± 0.20 | **< 1e−16** | **< 1e−16** | **2.4e-07** |
| | | pLDDT | 60.29±14.11 | 60.14±13.66 | 63.77±13.53 | 66.21±12.57 | **< 1e−16** | **< 1e−16** | **< 1e−16** |
